# Conflicting evolutionary histories of the mitochondrial and nuclear genomes in New World *Myotis* bats

**DOI:** 10.1101/112581

**Authors:** Roy N. Platt, Brant C. Faircloth, Kevin A.M. Sullivan, Troy Kieran, Travis C. Glenn, Michael W. Vandewege, Thomas E. Lee, Robert J. Baker, Richard D. Stevens, David A. Ray

## Abstract

The rapid diversification of *Myotis* bats into more than 100 species is one of the most extensive mammalian radiations available for study. Efforts to understand relationships within *Myotis* have primarily utilized mitochondrial markers and trees inferred from nuclear markers lacked resolution. Our current understanding of relationships within *Myotis* is therefore biased towards a set of phylogenetic markers that may not reflect the history of the nuclear genome. To resolve this, we sequenced the full mitochondrial genomes of 37 representative *Myotis,* primarily from the New World, in conjunction with targeted sequencing of 3,648 ultraconserved elements (UCEs). We inferred the phylogeny and explored the effects of concatenation and summary phylogenetic methods, as well as combinations of markers based on informativeness or levels of missing data, on our results. Of the 294 phylogenies generated from the nuclear UCE data, all are significantly different from phylogenies inferred using mitochondrial genomes. Even within the nuclear data, quartet frequencies indicate that around half of all UCE loci conflict with the estimated species tree. Several factors can drive such conflict, including incomplete lineage sorting, introgressive hybridization, or even phylogenetic error. Despite the degree of discordance between nuclear UCE loci and the mitochondrial genome and among UCE loci themselves, the most common nuclear topology is recovered in one quarter of all analyses with strong nodal support. Based on these results, we re-examine the evolutionary history of *Myotis* to better understand the phenomena driving their unique nuclear, mitochondrial, and biogeographic histories.

---

The bat genus *Myotis* (Order Chiroptera, Family Vespertilionidae) comprises more than 100 species that originated during the last 10-15 million years (Stadelmann et al. 2007), making it one of the most successful, extant, mammalian radiations. *Myotis* are distributed worldwide, excluding polar regions, and generally share a similar ecological niche: aerial insectivory. *Myotis* species often exhibit little morphological differentiation and, as a result, the rate of cryptic speciation within the genus is thought to be high. For example, specimens identified as *M. nigricans* and *M. albescens* form multiple paraphyletic lineages distributed throughout the phylogeny of Neotropical *Myotis* (Larsen et al. 2012). Confounding matters, the morphological variation that exists is often a poor indicator of species-level relationships. Early classifications of *Myotis* identified three major morphotypes (Findley 1972). Subsequent phylogenetic analyses of the mitochondrial cytochrome-b (*cytb*) gene demonstrated paraphyletic origins of each morphotype, suggesting frequent convergent evolution (Ruedi et al. 2001). These same analyses demonstrated that geography was a better predictor of phylogenetic relationship than morphology (Ruedi et al. 2001, Stadelmann et al. 2007).

The ability of mitochondrial markers to resolve a well-supported topology does not guarantee that the mitochondrial tree represents the species tree (for examples see Willis et al. 2014, Li, Gang et al. 2016, Leavitt et al. 2017). The lack of recombination and uniparental inheritance of the mitochondrion means that it is transmitted as a single genetic unit that is susceptible to evolutionary processes that may cause its history to diverge from the history of the species (Edwards et al. 2009). The most widely accepted phylogenies of *Myotis* rely heavily on mitochondrial data and even phylogenies containing nuclear data demonstrate an over reliance on mitochondrial markers for resolution, likely swamping out potentially conflicting signals from the nuclear genome. For example alignments of the nuclear *RAG2* and mitochondrial *cytb* contained 162 and 560 variable characters, respectively (Stadelmann et al. 2007). Despite these concerns, we find ourselves in a situation where our understanding of evolutionary relationships, our basis for conservation strategies, and our biogeographic hypotheses are all founded, almost entirely, on mitochondrial phylogenies.

Targeted sequencing of ultraconserved elements (UCEs; Faircloth et al. 2012) to collect sequence data from thousands of loci across the nuclear genome has resolved a number of difficult phylogenetic problems (for examples see Faircloth et al. 2012, McCormack et al. 2013, Green et al. 2014, Faircloth et al. 2015, McGee et al. 2016). Here we used UCE baits to collect ~1.4 Mbp from ≥3,600 nuclear loci in addition to random shotgun sequencing to collect full mitochondrial genomes in 37 taxa, primarily representing New World (NW) *Myotis.* Rather than analyzing each sequence data set once, or a handful of times, we chose to use a range of sampling, partitioning, and inference methods to fully explore phylogenetic tree space of the nuclear and mitochondrial genomes. We recovered 294 trees representing 175 distinct topologies for the nuclear UCE data and 28 trees representing 14 distinct mitochondrial topologies. Our results show that, despite the range of trees recovered from each marker type, nuclear and mitochondrial markers occupy distinct regions in tree space. Given that the nuclear and mitochondrial trees are distinct from one another it is necessary to reevaluate hypotheses made based solely on the mitochondrial phylogeny.

## Results

We used targeted sequencing of UCEs in 37 individuals (Table 1) to collect sequence data from 3,648 nuclear loci which we assembled into concatenated alignments as large as 1.37 Mb. In addition, we assembled mitochondrial genomes for most taxa from random shotgun sequencing data. We then used the data to infer the phylogenetic history of New World *Myotis* using a range of locus sampling strategies, alignment partitioning methods and phylogenetic inference methods.

**Table 1.** Specimens examined. Collection abbreviations: Museum of Southwestern Biology (MSB), Museum of Vertebrate Zoology (MVZ), Natural History Museum of Geneva (MHNG), Pontificia Universidad Catolica del Ecuador Museo de Zoologia (QCAZ), Texas Tech University Natural Science Research Laboratory (TK), University of Alaska Museum of the North (UAM), University of Michigan Museum of Zoology (UMMZ).

| Species | Name herein | Museum identification num. | Num. UCE Loci | UCE contig accession | Mitochondrial genome accession |
| --- | --- | --- | --- | --- | --- |
| *Eptesicus fuscus* | *fuscus*<sup>G</sup> | GCA_000308155.1 | 2,467 | Pending | KF111725.1 |
| *Eptesicus fuscus* | *fuscus* | TK 178736 | 2,849 | Pending | MF143474 |
| *Myotis albescens* | *albescens*<sup>1</sup> | RDS 7889 | 2,764 | Pending | MF143471 |
| *Myotis albescens* | *albescens*<sup>2</sup> | QCAZ 9157 | 1,185 | Pending | MF143497 |
| *Myotis albescens* | *albescens*<sup>3</sup> | TK 61766 | 2,872 | Pending | MF143470 |
| *Myotis ater* | *ater* | M4430 | 2,774 | Pending | MF143493 |
| *Myotis auriculus* | *auriculus* | MSB 40883 | 2,229 | Pending | MF143486 |
| *Myotis brandtii* | *brandtii*<sup>G</sup> | GCA_000412655.1 | 2,446 | Pending | KT210199.1 |
| *Myotis californicus* | *californicus* | UMMZ 175828 | 2,948 | Pending | MF143469 |
| *Myotis davidii* | *davidii*<sup>G</sup> | GCA_000327345.1 | 2,450 | Pending | KM233172.1 |
| *Myotis diminutus* | *diminutus* | QCAZ 9168 | 3,078 | Pending | MF143481 |
| *Myotis dominicensis* | *dominicensis* | TK 15624 | 2,576 | Pending | MF143467 |
| *Myotis evotis* | *evotis* | MSB 47323 | 2,586 | Pending | MF143468 |
| *Myotis fortidens* | *fortidens* | MSB 54941 | 2,791 | Pending | MF143483 |
| *Myotis horsfieldii* | *horsfieldii* | MHNG 1926.039 | 3,017 | Pending | MF143494 |
| *Myotis keaysi* | *keaysi* | TK 13525 | 3,195 | Pending | MF143477 |
| *Myotis keenii* | *keenii* | UAM 113849 | 2,723 | Pending | MF143472 |
| *Myotis leibii* | *leibii* | TK 24872 | 3,119 | Pending | MF143488 |
| *Myotis levis* | *levis* | RDS 7781 | 2,538 | Pending | MF143482 |
| *Myotis lucifugus* | *lucifugus*<sup>G</sup> | GCA_000147115.1 | 2,429 | Pending | KP273591.1 |
| *Myotis lucifugus* | *lucifugus* | MSB 46679 | 2,736 | Pending | MF143491 |
| *Myotis melanorhinus* | *melanorhinus* | TK 193888 | 3,177 | Pending | MF143489 |
| *Myotis nigricans* | *nigricans*<sup>1</sup> | QCAZ 9601 | 2,854 | Pending | MF143495 |
| *Myotis nigricans* | *nigricans*<sup>2</sup> | TK 194145 | 3,159 | Pending | MF143484 |
| *Myotis nyctor* | *nyctor* | TK 151413 | 856 | Pending | MF143498 |
| *Myotis occultus* | *occultus* | MSB 121995 | 2,957 | Pending | MF143490 |
| *Myotis oxyotus* | *oxyotus* | UMMZ RCO1013 | 3,106 | Pending | MF143479 |
| *Myotis riparius* | *riparius*<sup>1</sup> | TK 22688 | 2,924 | Pending | MF143473 |
| *Myotis riparius* | *riparius*<sup>2</sup> | TK 101723 | 2,990 | Pending | MF143475 |
| *Myotis riparius* | *riparius*<sup>3</sup> | TK 145199 | 2,890 | Pending | MF143480 |
| *Myotis ruber* | *ruber* | MVZ 185692 | 2,757 | Pending | MF143478 |
| *Myotis septentrionalis* | *septentrionalis* | TK 194420 | 2,916 | Pending | MF143487 |
| *Myotis thysanodes* | *thysanodes* | 07LEP | 2,821 | Pending | MF143492 |
| *Myotis velifer* | *velifer* | MSB 70877 | 2,704 | Pending | MF143499 |
| *Myotis vivesi* | *vivesi* | MSB 42658 | 2,469 | Pending | MF143476 |
| *Myotis volans* | *volans* | MSB 40886 | 2,819 | Pending | MF143496 |
| *Myotis yumanensis* | *yumanensis* | TK 194144 | 2,589 | Pending | MF143485 |

### Mitochondrial assembly, alignment, and phylogenetic inference

We generated an average of 5,498,440 paired reads per sample (min = 2,2370,68, max = 9,961,348) reads per sample of random DNA sequence data for mitochondrial genome assembly. Assemblies varied in quality. Some were almost entirely complete while others were missing small regions. We were not able to assemble through the control region for any of the samples. We found three premature stop codons in mtDNA protein coding genes. Subsequent manual alignment and validation suggested that these regions were miscalled by MitoBim, and we corrected the errors prior to analysis. Sequence coverage of the mitochondrial genomes averaged 58x (range >1× - 297×).

The thirty-seven mitochondrial protein coding, rRNA and tRNA genes were concatenated into an alignment of 15,520 bp containing 5,007 informative characters. rRNA and tRNA genes were concatenated into an alignment containing 4,157 bp containing 862 parsimony informative characters. Protein coding genes were concatenated into an alignment containing 11,363 bp or 3770 amino acids (aa) and containing 4,145 or 509 parsimony informative characters, respectively. For the alignment containing all protein coding, rRNA and tRNA genes, 30 samples were ≥ 90% complete, and alignments for five samples were 68-84% complete. Only 21% and 50% of nucleotide positions were present in the *M. albescens*^3^ (TK 61766) and *M. levis* alignments, respectively.

When considering alignment type, partitioning strategy, and phylogenetic inference method we analyzed the mitochondrial data 28 ways as described in the Methods. These analyses recovered 14 distinct topologies, the most common of which was recovered 6 times, all by analysis of the translated protein coding genes (Figure 1A). Despite being the most common topology, bipartition frequencies and clade probability values were lower than nucleotide based datasets, as would be expected with more slowly evolving amino acid sequences. Analysis of RNA coding genes recovered two topologies depending on the method of phylogenetic inference. The remainder of the analyses recovered various trees without any pattern relating to partitioning scheme or inference method. A majority rule consensus tree for all 28 analyses is shown in Figure 1C. The mitochondrial consensus tree reflects previously reported mitochondrial phylogenies for *Myotis.* The New World *Myotis* are generally monophyletic splitting into Nearctic and Neotropical clades with the Old World taxon, *M. brandtii,* sister to the Neotropical clade. Ambiguity, represented by polytomies, is found only in terminal relationships.

**Figure 1.**
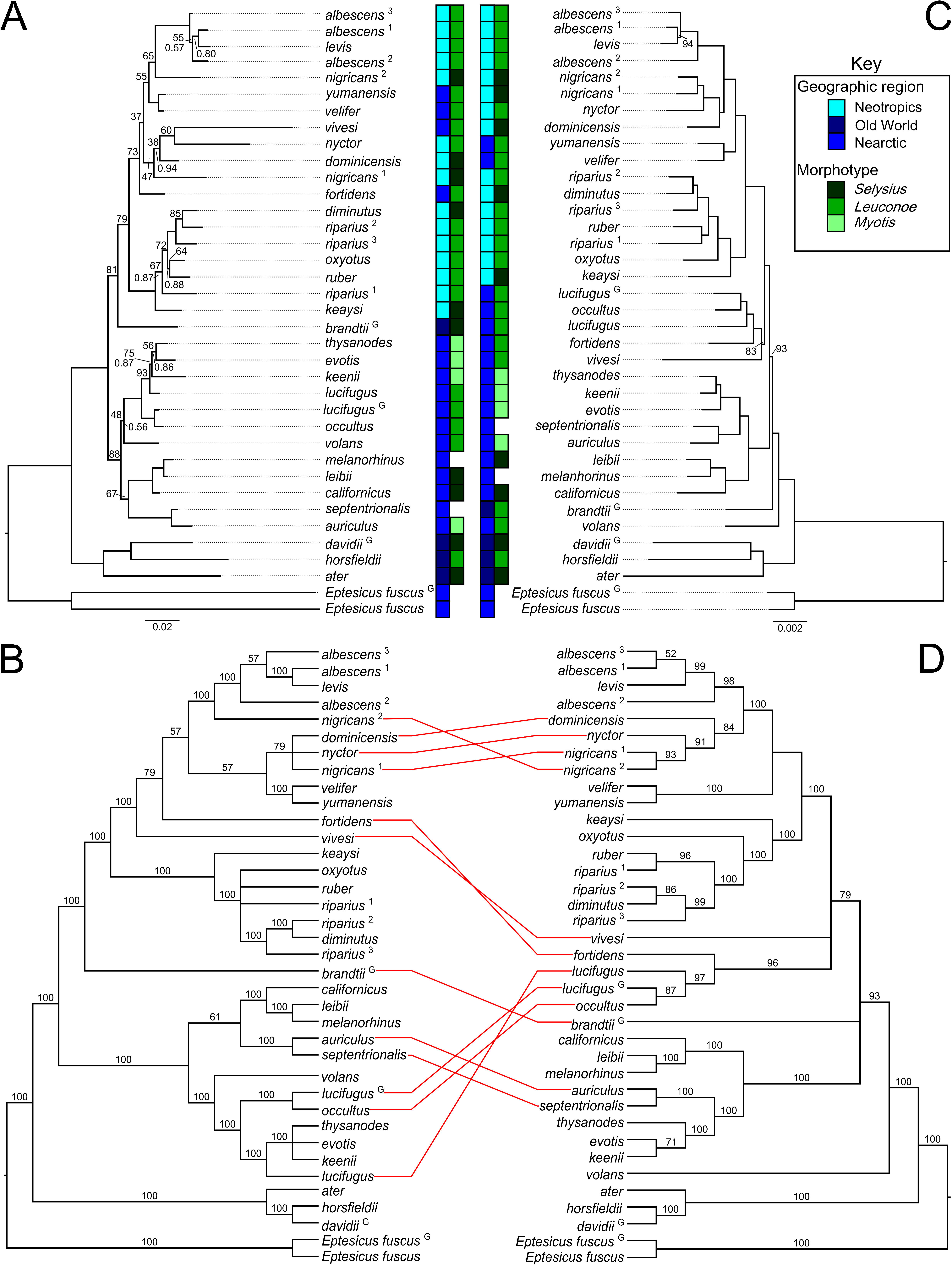
Comparison of nuclear and mitochondrial phylogenetic trees of *Myotis.* The most common mitochondrial (A) and nuclear (B) topologies. The mitochondrial topology arises from Bayesian and maximum likelihood analysis of the amino acid sequences portioned based on the PartitonFinder recommendations. The nuclear topology is from the 55% complete data matrix partitioned based on the recommendations of PartitionFinder. For clarity, bootstrap values greater than 95 and clade probability values greater than 0.95 have been omitted from the nuclear consensus tree. Majority-rule consensus trees from 28 mitochondrial (C) and 294 nuclear (D) topologies. Values above the branches in the consensus trees (C and D) are bipartition frequencies for that clade across all nuclear or mitochondrial topologies. Conflicting tips between data types (consensus nuclear vs. consensus mitochondrial) are indicated with red lines between the topologies. Biogeographic regions are color coded, as are subgeneric classifications based on morphotypes, as defined by Findley (1972). Species with more than one sample are designated with a superscript that is referenced in Table 1. Specimens derived from whole genome alignments are designated with a superscript “G”.

### UCE assembly and alignment, and phylogenetic inference

We averaged 3.29 million reads per sample after demultiplexing reads from the UCE-enriched, sequencing pool. These reads were assembled into an average of 5,778 contigs per sample (min = 1,562 *M. nyctor,* max = 11,784 *M. nigricans*^3^). Recovery rates for UCE loci varied across taxa. Of the 5,500 loci in the Amniote probe set, we successfully recovered 3,898 UCE loci, 3,648 loci from five or more samples and 212 loci in all 37 samples (Table 2). On average, 3,332 UCE loci were recovered per sample, ranging from 1,106 (*M. nyctor*) to 4,008 (*M. keaysi*). Repetitive sequences, identified via RepeatMasker searches, were minimal, occupying less than 0.02% of sites across all UCE alignments.

**Table 2.** General alignment information. For a subset of analyses a series of alignments were generated based on the number of taxa per locus. Thirty-seven taxa were examined so an alignment with all 37 taxa was considered 100% complete. Parsimony-informative characters make up a small portion of the total alignment. The number of partitions in the combined partitioning scheme was calculated with PartitionFinder.

| **Min. num. of taxa** | **Percent of**<br>**taxa per**<br>**locus** | **Number of Loci** | **Parsimony-**<br>**informative**<br>**characters** | **Variable**<br>**uninformative** | **Alignment**<br>**length** | **Optimum**<br>**Partitions** |
| --- | --- | --- | --- | --- | --- | --- |
| 5 | 15 | 3,648 | 38,718 | 62,588 | 1,377,262 | 31 |
| 9 | 25 | 3,379 | 37,894 | 57,672 | 1,260,248 | 37 |
| 12 | 35 | 3,232 | 37,259 | 56,453 | 1,227,093 | 34 |
| 16 | 45 | 3,064 | 36,539 | 54,782 | 1,187,492 | 31 |
| 20 | 55 | 2,890 | 35,284 | 53,200 | 1,144,471 | 27 |
| 24 | 65 | 2,668 | 34,373 | 51,148 | 1,091,620 | 27 |
| 27 | 75 | 2,481 | 33,031 | 48,732 | 1,041,099 | 20 |
| 31 | 85 | 2,034 | 29,179 | 41,711 | 903,903 | 16 |
| 35 | 95 | 1,193 | 18,288 | 24,189 | 575,321 | 15 |
| 37 | 100 | 212 | 3,480 | 4,778 | 112,125 | 6 |

In all, 294 different combinations of alignment, partitioning, and inference method were used to identify 175 unique nuclear topologies. The most common tree was recovered 45 times (Figure 1B). All nodes were highly-supported based on clade probability values and all but three nodes were highly supported with maximum likelihood bootstrap replicates. A consensus tree (Figure 1D) of the 294 nuclear topologies resolves New World *Myotis* (and *M. brandtii*) as a single polytomy that excludes *M. volans* and represents a departure from previous mitochondrial phylogenies, including the one recovered herein. Comparison of the mitochondrial and nuclear consensus trees ((Figures 1C and D) reveals substantial differences in the relationships within the genus.

As noted above, we were concerned that phylogenetic error, sampling bias or other factors may be driving the conflict between the mitochondrial and nuclear trees. We started by investigating the effects of matrix composition (or completeness) on phylogenetic inference using UCE loci by generating 10 alignments having varying levels of matrix completeness. Nine alignments were composed of loci with 15 to 95% (at 10% intervals) of taxa. The tenth alignment was composed of all loci which were recovered in 100% of specimens examined. For example, in the ‘15%’ matrix, all loci that were represented in at least 15% of species (n_taxa_ ≥ 5) were included in the alignment (n_loci_ = 3,648). In the ‘100%’ matrix, only loci which were identified in all species (n_taxa_ = 37) were included (n_loci_ = 212). Alignments were partitioned using three schemes: unpartitioned, individually partitioned by locus, or combined partitions. Alignment lengths, numbers of informative characters and number of partitions identified by PartitionFinder are available in Table 2. The result was 10 alignments that were each partitioned three ways. These were analyzed with Bayesian, maximum likelihood, and coalescent methods using RaxML, ExaBayes, ASTRID, ASTRAL-II and SVDquartets as described in the Methods. Bootstrap topologies stabilized in each alignment and partition combination within 150 replicates and all Bayesian runs converged in less than ten thousand generations. Unfortunately, computational limits forced us to abandon the individually partitioned, Bayesian analyses. In general, the same alignment produced the same topology regardless of inference method or partitioning scheme with the only exception being the terminal relationships of the *M. levis/M. albescens* clade in the combined vs. unpartitioned Bayesian analysis of the 100% complete data matrix.

Trees were also generated from data matrices incorporating UCE loci of differing lengths (Hosner et al. 2016). All 3,648 loci were grouped into ten bins based on locus length so that the first bin contained the 365 shortest loci, the second bin contained the 366^th^ to the 731^st^, and so on. The number of informative characters per bin ranged from 1,115 to 6,995 and the number of informative characters was correlated with average locus length (Supplemental Figure 1). On average, only 2.6% of characters in each bin were parsimony-informative (Supplemental Figure 1). Each of the ten length-based alignments recovered slightly different topologies. Terminal relationships were generally stable across analyses with the majority of differences between topologies found in the early bifurcations of the ingroup (*Myotis*).

From the above results, we observed that, in general, longer alignments produced well resolved topologies with significant nodal support regardless of the phylogenetic method or partitioning scheme used. Analyses of smaller data sets were more likely to recover unique topologies.

Given the goal of fully exploring nuclear tree space and generating as many reasonable nuclear UCE based topologies for comparison with the mitochondrial tree, we decided to randomly sample a limited number of nuclear UCE loci to create small alignments that would be more likely to result in unique topologies. We therefore randomly subsampled UCE loci to create 100 unique data sets of 365 loci. Loci were concatenated in each replicate data set and analyzed using maximum likelihood in RAxML. Of the 100 alignments analyzed, 80 unique topologies were generated (mean Robinson-Foulds distance = 4.3).

In addition to concatenation methods we explored phylogenetic tree space of the UCE data set using species tree methods. Normalized quartet scores from ASTRAL-II (Mirarab et al. 2015) analyses were consistent among all analyses, with scores ranging from 0.540 to 0.553, and the number of induced quartet gene trees ranging from 7,745,739 (100% complete 212 gene trees) and 63,042,410 (15% 3648 loci). SVDquartets (Chifman et al. 2014) sampled all 66,045 quartets. On average, the total weight of incompatible quartets was 2.84%. Similar to the concatenated analysis, we inferred coalescent-based species from the same 100 subsamples of 365 loci described above. Despite being generated from the same underlying data, summary and concatenation methods only recovered the same tree in 1 of 100 attempts.

Finally, we used weighted and unweighted statistical binning to combine individual gene trees into supergenes, estimate the supergene phylogeny, and then infer the species tree from the supergene trees. The 3,648 loci were combined into 528 binned loci with 480 bins containing seven loci each and 48 bins containing six loci each. Binning methods increases the normalized quartet scores from an average of 0.547 with gene trees to 0.672 for the binned-unweighted and 0.673 for the binned-weighted supergene trees. Given the relatively even distribution of loci into bins the negligible difference in quartet/species tree discordance between the unweighted and weighted supergene trees is expected. Binning methods have been shown to recover incorrect species trees under coalescent methods by altering gene tree frequencies. In addition, it is possible that support for incorrect trees can actually increase with the number of input gene trees (Liu et al. 2015). With these concerns in mind, both binning methods recovered the same topology which was the most common nuclear topology observed across all analyses (Figure 1C).

### Topological comparisons

All mitochondrial and nuclear trees are available in Supplemental File 1. When visualizing all topologies in tree space, nuclear trees co-localized and were distinct from mitochondrial topologies (Figure 2A). Comparisons of Robinson-Foulds symmetrical differences show 26 symmetrical differences between the most similar nuclear and mitochondrial trees (Figure 2B). For comparison only 9 of 43,070 and 0 of 378 pairwise nuclear vs. nuclear and mitochondrial vs. mitochondrial tree comparisons, Figures 2C and D, respectively, exhibited more than 26 symmetrical differences.

**Figure 2.**
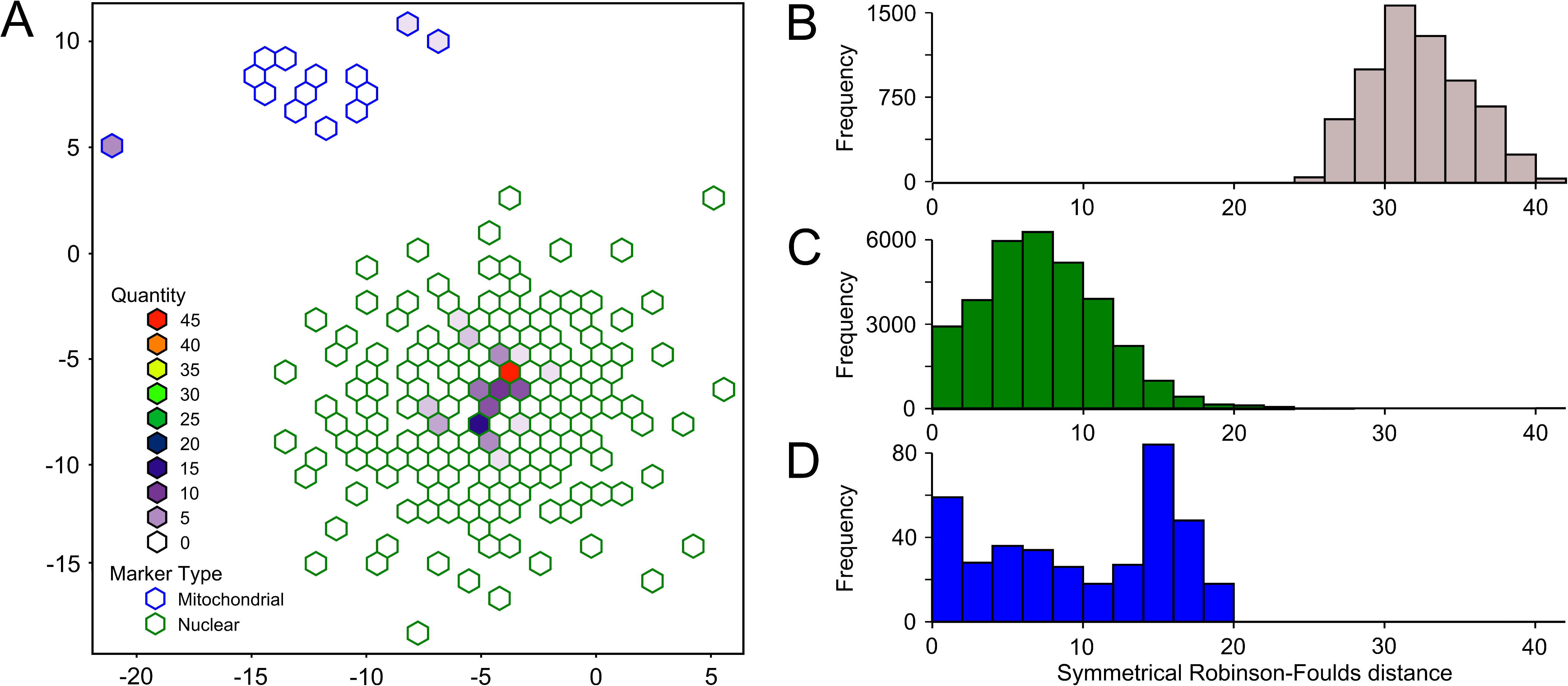
Comparison of mitochondrial and nuclear topologies. (A) All trees recovered from mitochondrial and nuclear analyses were visualized in tree space using multidimensional scaling of Robinson-Foulds distances. Nuclear trees (green outlines) were clustered separately from mitochondrial trees (blue outlines). Robinson-Foulds distances between nuclear and mitochondrial trees (B), nuclear vs. nuclear trees (C) and mitochondrial vs. mitochondrial trees (D) indicate the overall distinctions between the nuclear and mitochondrial topologies.

Of the 45 analyses that recovered the most frequently observed topology (Figure 1D), 38 were Bayesian and RAxML searches that varied by matrix completeness and partitioning scheme. The fact that these analyses recovered the same topology is expected given that they are not independent. For example, a RAxML analysis of the 25% complete data set uses 1.26 Mb of the 1.38 Mb of data present in the 15% complete matrix, sharing 91% of the data. Analyses that directly varied the alignments and/or sampled less data (e.g. randomly sampling loci) were more likely to generate unique topologies than the nested analyses described above. Of the 200 analyses that randomly sampled UCE loci 164 unique topologies were observed. This implies that when analyses of large data sets produce well-resolved trees with significant nodal support, sampling smaller portions of the data, may provide a mechanism for creating phylogenetic uncertainty not represented by typical tree scoring metrics.

## Discussion

We analyzed 3,648 UCE loci and mitochondrial genomes of 37 *Myotis* bats and outgroups. The mitochondrial and nuclear UCE phylogenies recovered distinct topologies (Figure 1), whether comparing the most commonly recovered or consensus topologies. Previous work with UCE loci demonstrated that support for deep divergences varied based on the number of loci examined (McCormack et al. 2013). Further, bootstrap replicates and clade probability values can be inaccurate metrics of nodal support (Douady et al. 2003, Hedtke et al. 2006). We used varied the input data and phylogenetic parameters to produce a range of reasonable nuclear and mitochondrial topologies that may be more useful than overreliance on a tree resulting from a one or two analyses. Rather than rejecting concordance between the two data types from a single analysis we took steps to analyze both data sets to generate as many viable topologies as possible, possibly revealing hidden overlap between marker types that are masked by phylogenetic error.

We recovered 175 and 14 unique nuclear and mitochondrial topologies from 294 and 28 analyses using multiple methodologies, sampling strategies, and phylogenetic parameters. By varying our phylogenetic methods we show that recovering a consistent topology is difficult even when using the same marker type, thus making it difficult to determine the “best” topology. Rather than choosing single trees to represent the nuclear and mitochondrial data, we use consensus trees to represent the ambiguity that is present through all analyses along with example phylogenies conforming to the most common topology (Figure 1). Despite our efforts to identify as many plausible, distinct nuclear and mitochondrial trees as possible we were unable recover overlapping topologies between marker types suggesting that nuclear UCE loci and the mitochondrial genomes of *Myotis* have distinct evolutionary histories.

A comparison of topologies in tree space illustrates the differenced in trees generated for each data set (Figure 2A). Intra-marker discordance, as measured by differences in nuclear vs. nuclear or mitochondrial vs. mitochondrial topologies is low. Pairwise comparisons of nuclear topologies contain an average of 8.3 symmetrical differences, mitochondrial comparisons contain an average of 10.8 differences. By contrast, pairwise differences between nuclear and mitochondrial topologies contain an average of 33.3 symmetrical differences. The two most similar nuclear and mitochondrial topologies contain 26 symmetrical differences. Of the 43,072 pairwise comparisons of nuclear topologies, only 27 contain more than 26 symmetrical differences. Likewise of the 378 pairwise comparisons of mitochondrial topologies only 1 contains more than 26 symmetrical differences. These observations indicate that discordance between marker types (Figure 2B) is much greater than within marker type (Figure 2C and 2D). All but the most divergent nuclear topologies are more similar to each other than any nuclear vs. mitochondrial comparison.

Conflict between mitochondrial and nuclear data may be driven by error in phylogenetic estimation or may reflect genuine discordance between the two marker types (Degnan et al. 2009, Huang, Huateng et al. 2010). We relied on multiple tree-inference methods (e.g. summary vs. concatenation), manipulated phylogenetic parameters (e.g. partitioning strategy), and sampling criteria (e.g. loci sampled in all taxa) to minimize the impacts of phylogenetic error on the data sets. In most cases, varying parameters or methodologies generated unique topologies with minor differences. In most cases unique topologies resulted from rearrangements of a few terminal taxa. For example, placement of *M. volans* and *M. brandtii* was often either sister to the remaining New World *Myotis* or as an early bifurcation between the Nearctic and Neotropical clades. *M. vivesi* was at times found as sister to the clade containing *M. lucifugus, occultus* and *fortidens* or as sister to the clade containing the neotropical *Myotis.*

Summary methods failed to recover the most common nuclear topology (Figure 1D) except when loci were binned prior to gene tree estimation. Summary methods also tended to recover more unique topologies than concatenation methods when analyzing data from the same gene(s). For example, the random sample analyses recovered 80 unique topologies using concatenation (RAxML) and 89 unique topologies with summary methods (ASTRAL-II). This likely has to do with the limited number of informative characters per locus and by extension limited phylogenetic signal per gene tree (Supplemental Figure 1). In these instances, limited phylogenetic signal per gene would likely lead to increased opportunity for phylogenetic error in gene tree estimation. Further supporting this idea, binning of compatible UCE loci may have indirectly increased phylogenetic signal resulting in the same topology that many of the concatenation analyses recovered. No other summary/coalescent method recovered this topology.

Previous studies that used primarily mitochondrial data recovered effectively the same relationships among *Myotis* as our mitochondrial analyses (Ruedi et al. 2001, Stadelmann et al. 2007, Roehrs et al. 2010, Larsen et al. 2012, Ruedi et al. 2013, Haynie et al. 2016). Thus, we are confident that the mitochondrial phylogeny we recovered here, and by others, reflects the true mitochondrial tree. However, the mitochondrial topology may not adequately reflect the species history, particularly when considering the factors that cause incongruence between nuclear and mitochondrial gene trees. Causes of conflicting gene trees include horizontal transfer, gene duplication, introgressive hybridization, and incomplete lineage sorting. The rapid radiation of this clade as well as other biological factors suggests that some of these phenomena are more likely to have influenced the *Myotis* radiation than others.

Horizontal transfer of genes is thought to be rare in eukaryotes, but, vespertilionids in general (Thomas et al. 2011, Platt et al. 2014, Platt et al. 2016), and *Myotis* in particular (Pritham et al. 2007, Ray et al. 2007, Ray et al. 2008, Pagan et al. 2010), have experienced horizontal transfer of DNA transposons. These events would not be reflected in our phylogeny because repetitive sequences were removed prior to phylogenetic analyses. More generally, gene duplications could create conflicting signal among individual UCE markers (ex. comparing non-orthologous UCE loci), but the number of gene duplication events would have to be very high to impact enough of the 3,648 UCE loci to confound the mitochondrial and nuclear phylogenies. Further ruling out gene duplication events as the dominant cause of conflicting phylogenetic signal is the fact that such events are likely depressed in *Myotis* as evidenced by their smaller genome size (~2.2 Gb) and trend towards DNA loss (Kapusta et al. 2017) combined with low rates of paralogy in UCEs general (Derti et al. 2006).

Introgressive hybridization and reticulation could significantly influence the phylogenies of *Myotis* in a way that leads to conflicting signal between the nuclear and mitochondrial genomes (Sota 2002, Good et al. 2015). Hybridization in bats may be relatively common given their propensity to swarm at cave entrances for breeding purposes. In European *Myotis,* swarming has allowed for high degrees of hybridization between *M. brandtii, M. mystacinus, and M. alcathoe* (Bogdanowicz et al. 2012). Further, *M. evotis, thysanodes,* and *keeni* all experienced historical gene flow during their divergence (Carstens et al. 2010, Morales et al. 2016). We could also explain the differences between the mitochondrial and nuclear UCE phylogenies if *Myotis* experienced extensive incomplete lineage sorting during their radiation. Two factors influence the rate of lineage sorting, the fixation rate and the speciation rate (Hudson et al. 2002). Increasing the time to fixation and/or decreasing the amount of time between cladogenic events will increase the likelihood of incomplete lineage sorting. *Myotis* are generally long lived species (Dzeverin 2008) and underwent a rapid radiation between 5-10 MYA (Lack et al. 2010), suggesting that *Myotis* species are likely to experience higher levels of lineage sorting. The importance of these events - introgressive hybridization and incomplete lineage sorting - in driving the differences between the mitochondrial and nuclear phylogenies should be further explored.

### Evolutionary history of *Myotis*

Our understanding of relationships within *Myotis* is heavily biased by mitochondrial data because nuclear markers were harder to collect and produced fewer informative sites (Ruedi et al. 2001, Stadelmann et al. 2007, Lack et al. 2010, Roehrs et al. 2010, Larsen et al. 2012, Ruedi et al. 2013, Haynie et al. 2016) or nuclear phylogenies with large numbers of makers contained limited numbers of taxa (Platt et al. 2015). Our UCE-based results indicate that nuclear trees vary substantially from the mitochondrial tree. Given that the nuclear and mitochondrial trees are different, we find it necessary to re-evaluate *Myotis* in the context of the nuclear data.

Paraphyly of *M. nigricans* and *M. albescens* was inferred from previous mitochondrial phylogenies. Larsen et al. (2012) identified a minimum of four and potentially twelve lineages in *M. albescens* and *M. nigricans.* Our sampling included three *M. albescens* and two *M. nigricans,* compared to Larsen’s 17 and 29 samples. Despite different mitochondrial and nuclear topologies overall, our mitochondrial and nuclear phylogeny recovered the same paraphyletic clade of three *M. albescens* samples and *M. levis.* Close relationships between these taxa was found in previous work and is expected. More importantly we did not find that *M. albescens* was paraphyletic across much of neotropical *Myotis.* We also found that *M. nigricans* is monphlyletic in the nuclear tree, but paraphyletic in the mitochondrial tree. These results from *M. nigricans* and *M. albescens* are interesting but further inference is limited due to low sample sizes for these taxa.

The original subgeneric taxonomy of *Myotis* was based on three morphotypes that were later shown to be the result of convergent evolution (Ruedi et al. 2001). If lineage-sorting affected the mitochondrial phylogeny, it is possible that the morphotypes truly are monophyletic. However, superimposing the previous subgeneric/morphological classification onto the species tree shows an interspersed distribution of morphotypes throughout the most common and consensus nuclear topologies (Figures 1A and B). Many strongly supported terminal relationships link species with different morphotypes. Based on these results, it appears that the three major morphotypes in *Myotis* are indeed a result of convergent evolution, as suggested by previous work (Ruedi et al. 2001, Stadelmann et al. 2007) despite the conflicting mitochondrial and nuclear phylogenies recovered

Among the more dramatic differences between the nuclear and mitochondrial topologies is the placement of *M. volans* and *M. brandtii* as sister to all New World taxa by the nuclear data. Our mitochondrial analyses place *M. volans* within a Nearctic clade and *M. brandtii* directly in-between the Nearctic and Neotropical bifurcations (Figure 1A and C) as has been previously reported (Stadelmann et al. 2007). By contrast the consensus UCE tree (Figure 1D) places *M. brondtii,* a species distributed throughout the Old World, within a polytomy at the base of the New World *Myotis* radiation and *M. volans* sister to all New World *Myotis* (including *M. brandtii*). The placement of *M. brandtii* in the nuclear UCE consensus tree does not necessarily contradict the mitochondrial phylogeny since it is an unresolved polytomy. However, it is worth noting that the most commonly recovered nuclear topology (Figure 1B) places *M. brandtii* sister to all New World *Myotis* (excluding *M. volans*). This relationship would more closely affiliate *M. brandtii* with other Old World taxa. The placement of *M. volans* as sister to all New World taxa (including *M. brandtii*) in the most common nuclear tree is a significant departure from previous work and, at first glance, does not make as much sense in a biogeographic framework. *M. volans* is distributed across western and northwestern North America as far as far north as Alaska. *M. brandtii* is distributed across much of Northern Europe and into the extreme eastern regions of Siberia. The key may lie in understanding the biogeography and phylogenetics of a third species, *M. gracilis*, a species that we were unfortunately unable to include.

*M. gracilis*, along with *M. brandtii*, are the only two *Myotis* geographically distributed in the Old World, but phylogenetically affiliated with the New World *Myotis* (Stadelmann et al. 2007). If future work verifies the sister relationship between *M. brandtii* and *M. gracilis,* then we can envision a scenario where *M. gracilis, M. brandtii,* and *M. volans* are the result of cladegenic events that occurred during the transition of *Myotis* from the Old World to the New World. It is important to remember that this interpretation relies on a fairly dramatic departure from the currently accepted mitochondrial relationships of *M. volans* (represented here by a single sample) to other *Myotis* species and abandons the nuclear UCE consensus tree for the most frequently recovered UCE topology. In addition, our taxonomic sampling excludes *Myotis* species from the OW, Ethiopian clade, the sister clade to New World *Myotis* (Ruedi et al. 2001, Stadelmann et al. 2007, Lack et al. 2010, Ruedi et al. 2013). Taxa from the Ethiopian clade are needed to properly root the New World *Myotis* clade and understand its biogeographic origins. With these caveats in mind, the hypothesis presented here should be viewed as highly speculative. Increasing the number of *Myotis* lineages sampled will shed additional light on this hypothesis.

Other taxa with conflicting positions between marker types include *M. lucifugus* + *M. occultus, M. fortidens,* and *M. vivesi.* In general, these relationships are characterized by very short branches (Figure 1D) and are the most likely to be affected by incomplete lineage sorting or limited phylogenetic information. This could explain the strong support with the mitochondrial tree compared to the nuclear species tree, while allowing for a number of nuclear loci to disagree with the species tree, as well.

There are a number of monophyletic groups identified with nuclear data (Figure 1A) that share general morphologies. For example, all of the long-eared bats (*septentrionalis, auriculus, evotis, thysanodes* and *keenii*) represent a monophyletic group of higher elevation, forest-dwelling species that glean insects off of surfaces (Fitch et al. 1979, O’Farrell et al. 1980, Warner 1982, Manning et al. 1989, Caceres et al. 2000). The group represented by *fortidens, lucifugus* and *occultus* represent a relatively long-haired form of *Myotis.* While having a distinct dental formula, *fortidens* was historically described as a subspecies of M. *lucifugus* (Miller Jr et al. 1928) and *occultus* has alternately represented its own species or been considered a subspecies of *lucifugus* (Hollister 1909, Valdez et al. 1999, Piaggio et al. 2002). The clade consisting of *keaysi, oxyotus, ruber, riparius,* and *diminutus* represents a neotropical group of primarily woolly-haired bats (LaVal 1973). None of these relationships are monophyletic in the consensus or most common mitochondrial topologies. If the mitochondrial genome has been subjected to phenomena that obscure the true species tree then these species groups, along with their synapomorphic morphological features, can be reevaluated.

### Conclusions

Relationships within *Myotis,* which until now have relied heavily on mitochondrial data, served as the basis for species identification (Puechmaille et al. 2012), evolutionary hypotheses (Simões et al. 2007), and even conservation recommendations (Boyles et al. 2007). Previous studies using nuclear data have largely been uninformative or utilized too few samples to draw definitive conclusions. Trees estimated from ~3,650 nuclear loci and 295 phylogenetic analyses recovered 175 topologies, none of which are congruent with the mitochondrial phylogeny of *Myotis*. Conflict between the mitochondrial and nuclear trees as well as among individual nuclear loci suggest that the *Myotis* radiation may have been accompanied by high levels of incomplete lineage sorting and possible hybridization. Rather than placing emphasis on the mitochondrial tree, it may be more appropriate to consider it for what it really is: a single gene on par with a single UCE locus, albeit one with many more phylogenetically informative characters. If true, then the mitochondrial genome is as likely to reflect the true species tree as any single UCE locus chosen at random. Phenomena such as lineage sorting, reticulation, and introgression have likely influenced the genomes of *Myotis* and should be accounted for in subsequent work. It is possible that the *Myotis* radiation is more accurately reflected as a hard polytomy or a phylogenetic network rather than a strictly bifurcating phylogeny.

## Materials and Methods

### Taxon Selection

Taxa were selected to span the major phylogenetic break points with emphasis on the Nearctic and Neotropical bifurcation as recovered in previous mitochondrial phylogenies (Stadelmann et al. 2007, Ruedi et al. 2013). In addition, multiple individuals morphologically identified as *M. nigricans* and *M. albescens* were included to test for paraphyly as suggested by Larsen et al. (2012). Three Old World species of *Myotis* and the outgroup, *E. fuscus,* were included to root phylogenetic analyses. All field identifications were confirmed from voucher museum specimens. Information for all specimens examined is available in Table 1.

### Library preparation, sequencing, and processing

Genomic DNA was extracted from 33 samples using either a Qiagen DNEasy extraction kit or a phenol-chloroform/ethanol precipitation. DNA was fragmented using the Bioruptor UCD-300 sonication device (Diagenode, Denville, NJ, USA). Libraries were prepared using the Kapa Library Preparation Kit KR0453-v2.13 (Kapa Biosystems, Wilmington, MA, USA) following the manufacturer’s instructions with five minor modifications. First, we used half volume reactions. Second, subsequent to end repair, we added Sera-Mag Speedbeads (Thermo-Scientific, Waltham, MA, USA; prepared according to (Glenn et al. 2016)) at a ratio of 2.86:1 for end repair cleanup. Third, we ligated universal iTru y-yoke adapters (Glenn et al. 2016) onto the genomic DNA. Fourth, following adapter ligation, we performed one post-ligation cleanup followed by Dual-SPRI size selection using 55 μL of speedbead buffer (22.5mM PEG, 1M NaCI) and 25 μL of Speedbeads. Finally, we performed a PCR at 95 °C for 45 sec, then 14 cycles of 98 °C for 30 sec, 60 °C for 30 sec, 72 °C for 30sec, then 72 °C for a 5 minute final extension and a 12 °C hold using iTru5 and iTru7 primers to produce Illumina TruSeqHT compatible libraries (Glenn et al. 2016).

Libraries were quantified on a Qubit 2.0 (Life Technologies) and 83 ng from each library was added to create 5 pools of 6 or 7 libraries each. We then split the pools in two. One subsample was enriched for UCE loci, the other was not. UCE loci in the enriched library pools were captured using Tetrapods 5K version 1 baits from MYcroarray (Ann Arbor, MI, USA) following their MYbaits protocol v. 2.3.1 with overnight incubations (Faircloth et al. 2012). Enriched libraries were quantified with a Qubit and pooled with other unrelated samples prior to sequencing on an Illumina HiSeq 3000 to produce paired-end reads of ≤ 151 bases. The unenriched samples were sequenced on a separate run using a single lane of Illumina HiSeq 2500. All samples were demultiplexed with Illumina’s fastq2bcl software. Reads were quality filtered by removing any potential adapter sequence and trimming read ends once the average Phred quality over a four base window score dropped below 20 using the Fastx toolkit (Gordon et al. 2010).

### Assembly, annotation, and phylogenetic analysis of the mitochondrial genome

Raw reads from the unenriched libraries were used to generate mitochondrial genomes via MitoBim (Hahn et al. 2013). This program used MIRA (Chevreux et al. 1999) to map reads to a *M. brandtii* reference genome (Genbank accession number KT210199.1). Alternative methods of mitochondrial genome assembly were used when MitoBim assembly failed. These taxa include *M. albescens* (TK61766), *M. albescens* (TK 101723), *M. albescens* (RDS 7889), *M. fortidens,* M. keeni *M. melanorhinus, M. nigricans* (QCAZ 9601), *M. septentrionalis, M. simus, M. velifer,* and *M. volans*. For these samples, we first identified reads that were mitochondrial in origin using BLAST searches against the *M. brandtii* mitochondrial genome (KT210199.1). Those reads were assembled using Trinity v2.2.0 with the ‘–single’ option to treat reads as unpaired. For taxa where we used NCBI genome assemblies to recover UCE loci *in silico* mitochondrial genomes from genbank were used to in the mitochondrial analyses GenBank as follows: *M. brandtii* (KT210199.1), *E. fuscas* (KF111725.1), *M. lucifugas* (KP273591.1), and *M. davidii* (KM233172.1).

Once assembled, each mitogenome was annotated via MITOS (Bernt et al. 2013). Annotated genes were manually validated via BLAST to confirm sequence identity and length. Protein coding genes were checked for stop codons using EMBOSS’s transeq program (Rice et al. 2000). When a stop codon was found, we used the raw reads to verify the sequence. We used BWA v0.7.12 (Li, Heng et al. 2009) to align the reads to the Mitobim assembled mitogenome to verify base calls from Mitobim. The protein coding rRNA and tRNA genes from each assembly were aligned using MUSCLE and concatenated into three different alignments containing only protein coding genes, only rRNA and tRNA genes, or all protein coding and RNA genes. A fourth alignment of all protein coding genes was translated to amino acids and concatenated into a single alignment.

Several different partitioning schemes were examined for each of the mitochondrial alignments. Alignments were either partitioned by gene, by codon, or by gene type (rRNA and tRNA vs. protein coding). Genes were partitioned individually except in the instances where two genes overlapped. These regions were partitioned separately from the individual genes resulting in three partitions for the two genes: a partition for gene A, a partition for gene B, and a partition for the overlapping nucleotides of gene A and B.

### Assembly, and phylogenetic analysis of nuclear UCEs

Quality filtered raw sequence reads from the UCE-enriched libraries were assembled into contigs using the Trinity assembler (Grabherr et al. 2011) with a minimum kmer coverage of 2. We used Phyluce to identify those assembled contigs that were UCE loci. We also harvested UCE loci from *Eptesicus fuscus* (GCA_000308155.1), *Myotis brandtii* (GCA_000412655.1), *M. davidii* (GCA_000327345.1), and *M. lucifugus* (GCF_000147115.1) genome assemblies using the Phyluce package (Faircloth 2016). Once extracted from Trinity and genome assemblies, we aligned all UCE loci MAFFT (Katoh et al. 2002) and trimmed the aligned data with gBIocks (Castresana 2000). Repetitive sequences (i. e. transposable elements) in each alignment were identified with RepeatMasker and trimmed where found.

Each alignment was analyzed using three different partitioning schemes. Unpartitioned alignments were simply concatenated UCE loci treated as a single unit. These alignments are referred to herein as “unpartitioned”. Fully partitioned alignments were concatenated alignments mitochondrial genes that were partitioned by locus. These alignments are referred to herein as “locus partitioned”. Finally, PartitionFinder v1.1.1 (Lanfear et al. 2012) was used to combine individual loci into an optimal partitioning scheme. The combined partitioning schemes for each alignment were identified with PartitionFinder v1.1.1 (Lanfear et al. 2012). Rather than searching for best-fit substitution models for each locus or partition, the GTR+Γ model of sequence evolution was assigned to all loci (Darriba et al. 2015) except in the case of amino acid alignments where the MtMam model was used. Initial trees for PartitionFinder were generated using RAxML v7.4.1 (Stamatakis 2006) with linked branch lengths. Partitioning schemes were heuristically searched using the hcluster algorithm. Partitioning schemes were chosen using the Bayeisan information criterion.

Finally, each alignment and partitioning scheme was analyzed using Bayesian inference and maximum likelihood phylogenetic methods. Bayesian phylogenies were generated with the MPI version of ExaBayes (v1.4.1) using 4 independent runs of 4 chains each. ExaBayes runs were terminated after 10 million generations only if the average standard deviation of split frequencies was less than 0.01. The first 25% of samples were discarded after which every 1,000^th^ generation was sampled. Proper sampling, post burn-in was inspected via Tracer v1.6. (Rambaut et al. 2014) and effective sample sizes greater than 200 were considered acceptable. Posterior probability values greater than 95% were considered to be significant. RaxML (v8.1.3) was used to estimate and score the maximum likelihood phylogeny with the rapid bootstrapping option and 10,000 bootstrap replicates. We define strongly supported bipartitions as those present in 95-100% of bootstrap replicates and moderately supported bipartitions are present in 85-95% of bipartitions (Wiens et al. 2008). All mitochondrial analyses are described in Table 3.

*Alignment types*. – Different combinations of UCE loci were used to create unique matrices based on 1) locus distribution, 2) locus length, or 3) random locus sampling. The first set of alignments (locus distribution) was determined by the number of taxa represented in the UCE alignments (phyluce_align_get_only_loci_with_min_taxa; Faircloth 2016), or degree of completeness. Matrices were constructed using loci which were present in 100% (number of specimens = 37), 95% (n = 35), 85% (n = 31), 75% (n = 27), 65% (n = 24), 55% (n = 20), 45% (n = 16), 35% (n = 12), 25% (n = 9), and 15% (n = 5) of specimens examined. These 10 groups were non-exclusive, so a locus that was assembled in all specimens (100% complete) would also be included with loci present in only 55% of specimens. On the other hand, a locus found in only 55% of specimens would not be included in the 100% complete data set. Each set of UCE alignments was concatenated using phyluce_align_format_nexus_files_for_raxml and a nexus character block was created using the phyluce_align_format_nexus_files_for_raxml – charsets option. These datasets then served as the basis for downstream phylogenetic analyses. For example, when a partitioning methodology (discussed below) was tested, it was performed for each of the 100%, 95%, 85%, etc. alignments. In addition to partitioning schemes, the effect of missing data was examined using Bayesian and maximum likelihood methods.

The second alignment criteria combined UCE loci of similar sizes. Pervious coalescent analyses of UCE data showed that sub-sampling the most informative loci can result in different topologies (Meiklejohn et al. 2016). Rather than using coalescent based analyses we used concatenation of UCE loci to identify different topologies based on length. Under these assumptions, UCE loci were sorted into ten groups based on their length and the predicted correlation between length and number of informative characters was confirmed (Supplemental Figure 1). UCE loci in the same size cohort were combined into a single alignment, partitioned by locus, and analyzed with RAxML using the methods described below.

The final set of alignments was generated by random sampling of UCE loci. In large phylogenetic analyses, systematic error can result in highly supported, but incorrect topologies due to compounding of non-phylogenetic signal (Rodríguez-Ezpeleta et al. 2007). By randomly reducing the dataset and replicating the maximum likelihood analyses, we can reduce the potential effects of compounding error. Roughly 10% of the dataset, 365 loci, were randomly sampled, concatenated, and partitioned by locus to create 100 new alignments, which were then analyzed with RAxML using the methods described below.

*Partitioning strategies.* –Alignments were analyzed using three different partitioning schemes – single, locus, and combined- similar to the mitochondrial partitioning schemes described above. Unpartitioned alignments were simply concatenated UCE loci treated as a single genetic unit. Rather than searching for best-fit substitution models for each UCE locus or partition, the GTR+Γ model of sequence evolution was assigned to all loci (Darriba et al. 2015). Initial trees for PartitionFinder were generated using RAxML v7.4.1 (Stamatakis 2006) with linked branch lengths. Partitioning schemes were heuristically searched using the hcluster algorithm and the best scheme was chosen using the Bayesian information criterion.

*Inference methods.* –Phylogenetic trees were generated with three different phylogenetic inference methods across five different inference implementations including concatenation and summary tree methods.

Maximum likelihood trees were inferred for each alignment and partitioning combination using RAxML v8.1.3 (Aberer et al. 2014). The best scoring (lowest -InL) tree from each dataset was identified from 100 random starting trees and bootstrapped 100 times using the GTR+Γ in both cases. The autoMRE function in RAxML v8.1.3 was used to determine the need for additional bootstrap replicates beyond the initial 100 (Pattengale et al. 2009). A stopping criterion was set *a priori* if the weighted Robinson-Foulds distance was less than 5% in 95% of random permutations of computed bootstrap replicates (Pattengale et al. 2009). If necessary, an additional 100 bootstrap replicates were computed until the convergence stopping criteria were met. Finally, bipartition frequencies of bootstrap replicates were drawn onto the best scoring tree from the initial RAxML searches for each of the respective data sets. Alignments based on UCE length or randomly sampled were analyzed using slightly different methods. In both cases UCE loci were partitioned individually and nodal support was calculated using 100 bootstrap replicates using the RAxML fast-bootstrapping option. For all maximum likelihood analyses, we define strongly supported bipartitions as those present in 95-100% of bootstrap replicates and moderately supported bipartitions are present in 85-95% of bipartitions (Wiens et al. 2008).

Bayesian analyses were conducted using ExaBayes v1.4.1 (Aberer et al. 2014). For all Bayesian analyses four independent runs of four chains each were run in parallel for a minimum of one million generations sampling every thousandth generation and applying a GTR+Γ substitution model for each partition. After one million generations, analyses continued until the standard deviation of the split frequency between chains was less than 0.01. The “-M 3” option was used to reduce the memory footprint of all ExaBayes runs. Proper sampling, post burn-in, was inspected via Tracer v1.6. (Rambaut et al. 2014). Effective sample sizes greater than 200 were considered acceptable. Posterior probability values greater than 95% were considered significant. An extended majority rule consensus tree was created from all trees after the first 25% of trees were discarded using TreeAnnotator v1.7.0 (Rambaut et al. 2013) and parameter estimates across all runs were calculated with Tracer vl.6 (Rambaut et al. 2014).

Species trees were calculated from gene trees for individual UCE loci recovered in five or more taxa using the GTR+Γ substitution model and fast bootstrapping (−f a) option in RAxML and 1,000 bootstrap replicates. In general, gene trees were classified based on the degree of completeness (i.e. number of taxa represented) similar to the way we treated individuals as described above.

Species trees were estimated and bootstrapped using three different programs. ASTRAL-II v4.10 (Mirarab et al. 2015) was used to build a summary tree. Support values for bipartitions in the tree were generated from 100 bootstrap replicates using site as well as site and locus resampling (Seo 2008). Species trees were estimated from ASTRID v1.4 (Vachaspati et al. 2015) using bionj and bootstrapped for 100 replicates. SVDquartets (Chifman et al. 2014), as implemented in PAUP v4.0a150 (Swofford 2003), was used to estimate a species trees from a random subset of 200,000 quartets and 1,000 bootstrap replicates.

UCE loci are relatively short markers with few informative characters from which to build gene trees. Errors in gene tree estimation may reduce the accuracy of summary methods and phylogenetic inference (Liu et al. 2009, Leaché et al. 2011, DeGiorgio et al. 2014, Mirarab et al. 2016). We used weighted and unweighted statistical binning to combine gene into compatible supergenes, increasing the number of informative informative characters per “locus” and reducing the phylogenetic error (Mirarab et al. 2014, Bayzid et al. 2015). The gene trees used for the summary tree methods described above were used rather than re-estimating trees. Bifurcations supported by more than 50% of the bootstrap replicates were retained for each gene tree. Alignments from compatible trees were concatenated into a single supergene alignment. Trees for supergenes were estimated using RAxML. The best trees for each supergene, as defined by log likelihood score, were retained from 500 searches. Bipartition support was estimated from 500 bootstrap replicates. For all analyses, the GTR+Γ model of substitution was used and each gene in the supergene alignment was partitioned separately. The resulting supertrees were then used for species tree estimation using ASTRAL-II. For the unweighted analysis, all supertrees were included in the pool of trees. For the weighted analysis, supertrees were weighted according to the number of genes combined in the supergene alignment. For example, if a supergene was a composite of six genes, the supertree was present 6 times compared to a composite of five genes which would be represented only five times. Support for the weighted and unweighted species trees was estimated by site and site and locus re-sampling (Seo 2008) for 100 bootstrap replicates in ASTRAL-II.

### Topological comparisons–

Trees recovered from all analyses were compared to each other in order to quantify the differences between topologies. Branch lengths have different meanings based on the type of analysis. For example, ASTRAL-II branch lengths are representative of coalescent units. ASTRID doesn’t even calculate branch lengths. For accurate tree comparisons, branch lengths were stripped from all trees. Pairwise unweighted Robinson-Foulds distances were calculated among all trees. Robinson-Foulds distances were transformed into two dimensions using the stochastic CCA algorithm for nonlinear dimension reduction in TreeScaper v1.09 (Huang, Wen et al. 2016). Coordinates were then visualized in R using hexagonal binning in the hexbin library v1.27.1 (Lewin-Koh 2011). Nuclear and mitochondrial 50% majority rule consensus trees were generated with PAUP v4.0a150 (Swofford 2003).

## Acknowledgments

We would like to thank the following museums, collection managers, and collaborators for the tissue loans necessary to complete this work: Joseph Cook (Museum of Southwestern Biology), Museum of Vertebrate Zoology, Manuel Ruedi (Natural History Museum of Geneva), Santiago Burneo (Pontificia Universidad Catolica del Ecuador Museo de Zoologia), Heath Garner, Robert Bradley and Caleb Phillips (Texas Tech University Natural Science Research Laboratory), Link Olson (University of Alaska Museum of the North), Cody Thompson and Priscilla Tucker (University of Michigan Museum of Zoology). In addition, we would like to thank the Texas Tech HPCC (http://www.depts.ttu.edu/hpcc/) for providing the computational resources necessary to complete this project. This work was supported by the National Science Foundation, DEB-1355176. Additional support was provided by College of Arts and Sciences at Texas Tech University. All sequence data is available at NCBI’s Short Read Archive (SRP095250). UCE contig assemblies are available at (PENDING ACCESSION NUMBER). Supplemental files include: a nexus file containing all estimated species trees (supFile-all_trees_2017-06-07.nexus.txt), and tables listing each of the phylogenetic analyses. An archived file of individual UCE alignments in fasta format (uceLociAlignments.tgz), and an archived file containing all individual UCE gene trees in newick format (UceGeneTrees.tgz) are available through DRYAD (PENDING ACCESSION NUMBER).

**Sup Fig 1.** Phylogenetic content of UCE loci based on length. The length of a UCE locus is correlated with the number of phylogenetically informative characters. UCE loci were sorted by length and ten bins of alignments were created so that the shortest loci were combined into one alignment, the next shortest set of loci were combined … etc. Parsimony informative characters made up a minor part of each alignment.

**Sup File 1** - supFile-all_trees_2017-06-07.nexus.txt –Trees generated from all analyses in nexus format.

## Literature Cited

Aberer A. J., Kobert K., Stamatakis A. 2014. ExaBayes: Massively parallel Bayesian tree inference for the whole-genome era. Mol Biol Evol. 31:2553-2556.

Bayzid M. S., Mirarab S., Boussau B., Warnow T. 2015. Weighted statistical binning: Enabling statistically consistent genome-scale phylogenetic analyses. PLoS ONE. 10:e0129183.

Bernt M., Donath A., Jühling F., Externbrink F., Florentz C., Fritzsch G., Pütz J., Middendorf M., Stadler P. F. 2013. MITOS: Improved de novo metazoan mitochondrial genome annotation. Mol Phylogenet Evol. 69:313-319.

Bogdanowicz W., Piksa K., Tereba A. 2012. Hybridization hotspots at bat swarming sites. PLoS ONE. 7:e53334.

Boyles J. G., Storm J. J. 2007. The perils of picky eating: Dietary breadth is related to extinction risk in insectivorous bats. PLOS ONE. 2:e672.

Caceres M. C., Barclay R. M. R. 2000. Myotis septentrionalis. Mammalian Species:l-4.

Carstens B. C., Dewey T. A. 2010. Species delimitation using a combined coalescent and information-theoretic approach: An example from North American *Myotis* bats. Syst Biol. 59:400-414.

Castresana J. 2000. Selection of conserved blocks from multiple alignments for their use in phylogenetic analysis. Mol Biol Evol. 17:540-552.

Chevreux B., Wetter T., Suhai S. 1999. Genome sequence assembly using trace signals and additional sequence information. Proceedings of the German Conference on Bioinformatics (GCB).

Chifman J., Kubatko L. 2014. Quartet inference from SNP data under the coalescent model. Bioinformatics. 30:3317-3324.

Darriba D., Posada D. 2015. The impact of partitioning on phylogenomic accuracy. bioRxiv. https://doi.org/10.1101/023978.

DeGiorgio M., Degnan J. H. 2014. Robustness to divergence time underestimation when inferring species trees from estimated gene trees. Syst Biol. 63:66-82.

Degnan J. H., Rosenberg N. A. 2009. Gene tree discordance, phylogenetic inference and the multispecies coalescent. Trends Ecol Evol. 24:332-340.

Derti A., Roth F. P., Church G. M., Wu C. t. 2006. Mammalian ultraconserved elements are strongly depleted among segmental duplications and copy number variants. Nat Genet. 38:1216-1220.

Douady C. J., Delsuc F., Boucher Y., Doolittle W. F., Douzery E. J. P. 2003. Comparison of Bayesian and maximum likelihood bootstrap measures of phylogenetic reliability. Mol Biol Evol. 20:248-254.

Dzeverin I. 2008. The stasis and possible patterns of selection in evolution of a group of related species from the bat genus *Myotis* (Chiroptera, Vespertilionidae). J Mamm Evol. 15:123-142.

Edwards S., Bensch S. 2009. Looking forwards or looking backwards in avian phylogeography? A comment on Zink and Carrowclough 2008. Mol Ecol. 18:2930-2933.

Faircloth B. C. 2016. PHYLUCE is a software package for the analysis of conserved genomic loci. Bioinformatics. 32:786-788.

Faircloth B. C., Branstetter M. G., White N. D., Brady S. G. 2015. Target enrichment of ultraconserved elements from arthropods provides a genomic perspective on relationships among Hymenoptera. Mol Ecol Resour. 15:489-501.

Faircloth B. C., McCormack J. E., Crawford N. G., Harvey M. G., Brumfield R. T., Glenn T. C. 2012. Ultraconserved elements anchor thousands of genetic markers spanning multiple evolutionary timescales. Syst. Biol. 61:717-726.

Findley J. S. 1972. Phenetic relationships among bats of the genus *Myotis*. Syst Biol. 21:31-52.

Fitch J. H., Shump K. A. 1979. Myotis keenii. Mammalian Species Archive. 121:1-3.

Glenn T. C., Nilsen R., Kieran T. J., Finger J. W., Pierson T. W., Bentley K. E., Hoffberg S., Louha S., Garcia-De-Leon F. J., Angel del Rio PortillaM., Reed K., Anderson J. L., Meece J. K., Aggery S., Rekaya R., Alabady M., Belanger M., Winker K., Faircloth B. C. 2016. Adapterama I: Universal stubs and primers for thousands of dual-indexed Illumina libraries (iTru & iNext). bioRxiv. https://doi.org/10.1101/049114.

Good J. M., Vanderpool D., Keeble S., Bi K. 2015. Negligible nuclear introgression despite complete mitochondrial capture between two species of chipmunks. Evolution. 69:1961-1972.

Gordon A., Hannon G. 2010. Fastx-toolkit. http://hannonlab.cshl.edu/fastx_toolkit.

Grabherr M. G., Haas B. J., Yassour M., Levin J. Z., Thompson D. A., Amit I., Adiconis X., Fan L., Raychowdhury R., Zeng Q., Chen Z., Mauceli E., Hacohen N., Gnirke A., Rhind N., di Palma F., Birren B. W., Nusbaum C., Lindblad-Toh K., Friedman N., Regev A. 2011. Full-length transcriptome assembly from RNA-Seq data without a reference genome. Nat Biotech. 29:644-652.

Green R. E., Braun E. L., Armstrong J., Earl D., Nguyen N., Hickey G., Vandewege M. W., St. John J. A., Capella-Gutiérrez S., Castoe T. A., Kern C., Fujita M. K., Opazo J. C., Jurka J., Kojima K. K., Caballero J., Hubley R. M., Smit A. F., Platt R. N., Lavoie C. *A.*, Ramakodi M. P., Finger J. W., Suh A., Isberg S. R., Miles L., Chong A. Y., Jaratlerdsiri W., Gongora J., Moran C., Iriarte A., McCormack J., Burgess S. C., Edwards S. V., Lyons E., Williams C., Breen M., Howard J. T., Gresham C. R., Peterson D. G., Schmitz J., Pollock D. D., Haussler D., Triplett E. W., Zhang G., Irie N., Jarvis E. D., Brochu C. *A.*, Schmidt C. J., McCarthy F. M., Faircloth B. C., Hoffmann F. G., Glenn T. C., Gabaldón T., Paten B., Ray D. A. 2014. Three crocodilian genomes reveal ancestral patterns of evolution among archosaurs. Science. 346:1254449.

Hahn C., Bachmann L., Chevreux B. 2013. Reconstructing mitochondrial genomes directly from genomic next-generation sequencing reads—a baiting and iterative mapping approach. Nucleic Acids Res. 41:e129.

Haynie M. L., Tsuchiya M. T. N., Ospina-Garcés S. M., Arroyo-Cabrales J., Medellín R. *A.*, Polaco O. J., Maldonado J. E. 2016. Placement of the rediscovered *Myotis planiceps* (Chiroptera: Vespertilionidae) within the *Myotis* phylogeny. J Mammal. 97:701-712.

Hedtke S. M., Townsend T. M., Hillis D. M. 2006. Resolution of phylogenetic conflict in large data sets by increased taxon sampling. Syst Biol. 55:522-529.

Hollister N. 1909. Two new bats from the southwestern United States, publisher not identified.

Hosner P. *A.*, Faircloth B. C., Glenn T. C., Braun E. L., Kimball R. T. 2016. Avoiding missing data biases in phylogenomic inference: An empirical study in the landfowl (Aves: Galliformes). Mol Biol Evol. 33:1110-1125.

Huang H., He Q., Kubatko L. S., Knowles L. L. 2010. Sources of error inherent in species-tree estimation: Impact of mutational and coalescent effects on accuracy and implications for choosing among different methods. Syst Biol. 59:573-583.

Huang W., Zhou G., Marchand M., Ash J. R., Morris D., Van Dooren P., Brown J. M., Gallivan K. *A.*, Wilgenbusch J. C. 2016. TreeScaper: Visualizing and extracting phylogenetic signal from sets of trees. Mol Biol Evol.

Hudson R. R., Coyne J. *A.*, Huelsenbeck J. 2002. Mathematical consequences of the genealogical species concept. Evolution. 56:1557-1565.

Kapusta A., Suh A., Feschotte C. 2017. Dynamics of genome size evolution in birds and mammals. Proc Natl Acad Sci U S A. 114:E1460-E1469.

Katoh K., Misawa K., Kuma K. i., Miyata T. 2002. MAFFT: a novel method for rapid multiple sequence alignment based on fast Fourier transform. Nucleic Acids Res. 30:3059-3066.

Lack J. B., Roehrs Z. P., Stanley C. E., Ruedi M., Van Den Bussche R. A. 2010. Molecular phylogenetics of *Myotis* indicate familial-level divergence for the genus *Cistugo* (Chiroptera). J Mammal. 91:976-992.

Lanfear R., Calcott B., Ho S. Y. W., Guindon S. 2012. PartitionFinder: Combined selection of partitioning schemes and substitution models for phylogenetic analyses. Mol Biol Evol. 29:1695-1701.

Larsen R. J., Knapp M. C., Genoways H. H., Khan F. A. *A.*, Larsen P. *A.*, Wilson D. E., Baker R. J. 2012. Genetic diversity of neotropical *Myotis* (Chiroptera: Vespertilionidae) with an emphasis on South American species. PLoS ONE. 7:e46578.

LaVal R. K. 1973. A revision of the Neotropical bats of the genus *Myotis*. Natural History Museum, Los Angeles County.

Leaché A. D., Rannala B. 2011. The accuracy of species tree estimation under simulation: A comparison of methods. Syst Biol. 60:126-137.

Leavitt D. H., Marion A. B., Hollingsworth B. D., Reeder T. W. 2017. Multilocus phylogeny of alligator lizards (Elgaria, Anguidae): Testing mtDNA introgression as the source of discordant molecular phylogenetic hypotheses. Mol Phylogenet Evol.

Lewin-Koh N. 2011. Hexagon binning: an overview. ftp://ftp.naist.jp/pub/lang/R/CRAN/web/packages/hexbin/vignettes/hexagon_binning.pdf.

Li G., Davis B. W., Eizirik E., Murphy W. J. 2016. Phylogenomic evidence for ancient hybridization in the genomes of living cats (Felidae). Genome Res. 26:1-11.

Li H., Durbin R. 2009. Fast and accurate short read alignment with Burrows–Wheeler transform. Bioinformatics. 25:1754-1760.

Liu L., Edwards S. V. 2015. Comment on “Statistical binning enables an accurate coalescent-based estimation of the avian tree”. Science. 350:171-171.

Liu L., Yu L., Kubatko L., Pearl D. K., Edwards S. V. 2009. Coalescent methods for estimating phylogenetic trees. Mol Phylogenet Evol. 53:320-328.

Manning R. W., Jones J. K. 1989. Myotis evotis. Mammalian Species Archive. 329:1-5.

McCormack J. E., Harvey M. G., Faircloth B. C., Crawford N. G., Glenn T. C., Brumfield R. T. 2013. A Phylogeny of birds based on over 1,500 loci collected by target enrichment and high-throughput sequencing. PLoS ONE. 8:e54848.

McGee M. D., Faircloth B. C., Borstein S. R., Zheng J., Darrin Hulsey C., Wainwright P. C., Alfaro M. E. 2016. Replicated divergence in cichlid radiations mirrors a major vertebrate innovation. Proc R Soc Lond B Biol Sc. 283.

Meiklejohn K. A., Faircloth B. C., Glenn T. C., Kimball R. T., Braun E. L. 2016. Analysis of a rapid evolutionary radiation using Ultraconserved Elements: Evidence for a bias in some multispecies coalescent methods. Syst Biol. 65:612-627.

Miller Jr G. S., Allen G. 1928. The American bats of the genus *Myotis* and *Pixonyx*. Bulletin of the United States National Museum. 144:1-218.

Mirarab S., Bayzid M. S., Boussau B., Warnow T. 2014. Statistical binning enables an accurate coalescent-based estimation of the avian tree. Science. 346.

Mirarab S., Bayzid M. S., Warnow T. 2016. Evaluating summary methods for multilocus species tree estimation in the presence of incomplete lineage sorting. Syst Biol. 65:366-380.

Mirarab S., Warnow T. 2015. ASTRAL-II: Coalescent-based species tree estimation with many hundreds of taxa and thousands of genes. Bioinformatics. 31:i44-i52.

Morales A. E., Jackson N. D., Dewey T. A., O’Meara B. C., Carstens B. C. 2016. Speciation with gene flow in North American *Myotis* bats. Syst Biol.

O’Farrell M. J., Studier E. H. 1980. Myotis thysanodes. Mammalian Species. 137:1-5.

Pagan H. J. T., Smith J. D., Hubley R. M., Ray D. A. 2010. PiggyBac-ing on a Primate genome: Novel elements, recent activity and horizontal transfer. Genome Biol Evol. 2.

Pattengale N. D., Alipour M., Bininda-Emonds O. R. P., Moret B. M. E., Stamatakis A. 2009. How many bootstrap replicates are necessary? In: Batzoglou S editor. Research in Computational Molecular Biology: 13th Annual International Conference, RECOMB 2009, Tucson, AZ, USA, May 18-21, 2009.

Proceedings. Berlin, Heidelberg, Springer Berlin Heidelberg, p. 184-200.

Piaggio A. J., Valdez E. W., Bogan M. A., Spicer G. S. 2002. Systematics of *Myotis occultus* (Chiroptera: Vespertilionidae) inferred from sequences of two mitochondrial genes. J Mammal. 83:386-395.

Platt R. N., Mangum S. F., Ray D. A. 2016. Pinpointing the vesper bat transposon revolution using the Miniopterus natalensis genome. Mob DNA. 7:12.

Platt R. N., Vandewege M. W., Kern C., Schmidt C. J., Hoffmann F. G., Ray D. A. 2014. Large numbers of novel miRNAs originate from DNA transposons and are coincident with a large species radiation in bats. Mol Biol Evol. 31:1536-1545.

Platt R. N., Zhang Y., Witherspoon D. J., Xing J., Suh A., Keith M. S., Jorde L. B., Stevens R. D., Ray D. A. 2015. Targeted Capture of Phylogenetically Informative Ves SINE Insertions in Genus Myotis. Genome Biol Evol. 7:1664-1675.

Pritham E. J., Feschotte C. 2007. Massive amplification of rolling-circle transposons in the lineage of the bat *Myotis* lucifugus. Proc Natl Acad Sci U S A. 104:1895-1900.

Puechmaille S. J., Allegrini B., Boston E. S. M., Dubourg-Savage M.-J., Evin A., Knochel A., Le Bris Y., Lecoq V., Lemaire M., Rist D., Teeling E. C. 2012. Genetic analyses reveal further cryptic lineages within the *Myotis nottereri* species complex. Mammalian Biology. 77:224-228.

Rambaut A., Drummond A. 2013. TreeAnnotator v1. 7.0. http://beast.bio.ed.ac.uk.

Rambaut A., Suchard M., Xie D., Drummond A. 2014. Tracer v1.6. http://beast.bio.ed.ac.uk.

Ray D. *A.*, Feschotte C., Pagan H. J. T., Smith J. D., Pritham E. J., Arensburger P., Atkinson P. W., Craig N. L. 2008. Multiple waves of recent DNA transposon activity in the bat, *Myotis lucifugus*. Genome Res. 18:717-728.

Ray D. *A.*, Pagan H. J. T., Thompson M. L., Stevens R. D. 2007. Bats with hATs: Evidence for recent DNA transposon activity in genus *Myotis*. Mol Biol Evol. 24:632-639.

Rice P., Longden I., Bleasby A. 2000. EMBOSS: the European molecular biology open software suite. Elsevier Current Trends.

Rodríguez-Ezpeleta N., Brinkmann H., Roure B., Lartillot N., Lang B. F., Philippe H. 2007. Detecting and overcoming systematic errors in genome-scale phylogenies. Syst Biol. 56:389-399.

Roehrs Z. P., Lack J. B., Van Den Bussche R. A. 2010. Tribal phylogenetic relationships within Vespertilioninae (Chiroptera: Vespertilionidae) based on mitochondrial and nuclear sequence data. J Mammal. 91:1073-1092.

Ruedi M., Mayer F. 2001. Molecular systematics of bats of the genus *Myotis* (Vespertilionidae) suggests deterministic ecomorphological convergences. Mol Phylogenet Evol. 21:436-448.

Ruedi M., Stadelmann B., Gager Y., Douzery E. J. P., Francis C. M., Lin L.-K., Guillén-Servent A., Cibois A. 2013. Molecular phylogenetic reconstructions identify East Asia as the cradle for the evolution of the cosmopolitan genus *Myotis* (Mammalia, Chiroptera). Mol Phylogenet Evol. 69:437-449.

Seo T.-K. 2008. Calculating bootstrap probabilities of phylogeny using multilocus sequence data. Mol Biol Evol. 25:960-971.

Simôes B. F., Rebelo H., Lopes R. J., Alves P. C., Harris D. J. 2007. Patterns of genetic diversity within and between *Myotis d. daubentonii* and *M. d. nathalinae* derived from cytochrome *b* mtDNA sequence data. Acta Chiropt. 9:379-389.

Sota T. 2002. Radiation and reticulation: extensive introgressive hybridization in the carabid beetles *Ohomopterus* inferred from mitochondrial gene genealogy. Population Ecology. 44:0145-0156.

Stadelmann B., Lin L. K., Kunz T. H., Ruedi M. 2007. Molecular phylogeny of New World *Myotis* (Chiroptera, Vespertilionidae) inferred from mitochondrial and nuclear DNA genes. Mol Phylogenet Evol. 43:32-48.

Stamatakis A. 2006. RAxML-VI-HPC: maximum likelihood-based phylogenetic analyses with thousands of taxa and mixed models. Bioinformatics. 22:2688-2690.

Swofford D. L. 2003. PAUP*. Phylogenetic analysis using parsimony (* and other methods). Version 4.

Thomas J., Sorourian M., Ray D., Baker R. J., Pritham E. J. 2011. The limited distribution of Helitrons to vesper bats supports horizontal transfer. Gene. 474:52-58.

Vachaspati P., Warnow T. 2015. ASTRID: Accurate Species TRees from Internode Distances. BMC Genomics. 16:S3.

Valdez E. W., Choate J. R., Bogan M. A., Yates T. L. 1999. Taxonomic status of *Myotis occultus*. J Mammal. 80:545-552.

Warner R. M. 1982. Myotis auriculus. Mammalian Species. 191:1-3.

Wiens J. J., Kuczynski C. A., Smith S. *A.*, Mulcahy D. G., Sites J. J. W., Townsend T. M., Reeder T. W. 2008. Branch lengths, support, and congruence: Testing the phylogenomic approach with 20 nuclear loci in snakes. Syst Biol. 57:420-431.

Willis S. C., Farias I. P., Ortí G. 2014. Testing mitochondrial capture and deep coalescence in Amazonian cichlid fishes (Cichlidae: *Cichla*). Evolution. 68:256-268.

